# Evidence for the acquisition of a proteorhodopsin-like rhodopsin by a chrysophyte-infecting giant virus

**DOI:** 10.1101/2025.06.17.660233

**Authors:** Petra Byl, Christopher R. Schvarcz, Julie Thomy, Qian Li, Cori B. Williams, Kurt LaButti, Frederik Schulz, Kyle F. Edwards, Grieg F. Steward

## Abstract

Chrysophytes are nanoflagellate protists widespread in aquatic ecosystems with diverse trophic roles as primary producers and bacterivores. Molecular evidence suggests that chrysophytes are commonly infected by giant viruses but isolates of such virus-host systems have not been reported. Here, we describe the first cultivated chrysophyte-infecting virus, Chrysophyceae Clade H virus SA1 (ChrysoHV), isolated along with its phago-mixotrophic host alga from surface waters in the tropical North Pacific Ocean. The ChrysoHV capsid (290 ± 40 nm diameter) is associated with a loose, sac-like membrane that extends its effective diameter (720 ± 120 nm) and presents a long (1,200 ± 240 nm), thin (20 ± 2), flexible tail, a morphology unlike any virion yet described. The assembled genome is 1.19 Mbp. Phylogenetic analysis places ChrysoHV as the third cultivated member of the *Aliimimivirinae* subfamily in the *Mimiviridae* family of giant viruses. The ChrysoHV genome encodes two heliorhodopsins and one proteorhodopsin. Proteorhodopsins are well known light-driven proton pumps in bacteria but have not been previously reported in a viral genome. The predicted viral proteorhodopsin structure suggests it may not have a functional retinal binding site implying a light-independent function. The genome also encodes two ribosomal proteins and nine genes with closest known homologs in marine cyanobacteria, most annotated as encoding for proteins involved in nutrient uptake. This unusual virus could serve as a model system for exploring viral rhodopsin functions and its genome suggests that phagotrophic protists may serve as an intracellular market for gene exchange between infecting viruses and ingested bacterial prey.

**Importance:** Chrysophytes are abundant eukaryotic phytoplankton with trophic strategies ranging from photosynthesis to phagotrophy. They serve as models of mixotrophy among aquatic protists, but no chrysophyte-infecting viruses had been isolated, leaving a gap in experimental virus-host systems for a major class of protists. This study reports on the characterization of the first isolated chrysophyte-infecting virus, ChrysoHV. The virion morphology is unusual, having a loose membranous sac around a large capsid and a long filamentous tail. The genome contains genes for ribosomal proteins, a rarity in eukaryotic viruses, and multiple genes with homologs in common marine bacteria, one of which is a type of rhodopsin never before reported in a virus. We hypothesize that phago-mixotrophs, through infections and ingestion, may facilitate lateral gene exchange between eukaryote-infecting viruses and bacteria, entities that might not otherwise directly interact. The results expand the observed morphological diversity among viruses and the catalog of known virus genes.

## Introduction

Chrysophytes are single-celled or colonial protists in the class Chrysophyceae, and they are globally abundant in freshwater and marine ecosystems (1, 2). Chrysophytes play multiple ecosystem roles as primary producers and bacterivores, and include phago-mixotrophic forms that perform photosynthesis and ingest prey, strictly heterotrophic phagotrophs, and potentially non-phagotrophic phototrophs (3). In subtropical ocean gyre ecosystems, which span approximately a quarter of the global surface ocean (4), chrysophytes play an important role as predators of marine cyanobacteria (5, 6), influencing the community structure at the base of the food web. Isolates from Chrysophyceae Clade H have been shown to be voracious consumers of *Prochlorococcus* (7), a cyanobacterium that is the dominant primary producer in subtropical gyres and most abundant phototroph in the ocean (8).

There is also considerable evidence of interactions between chrysophytes and dsDNA viruses. Chrysophyte genomes can harbor tens to hundreds of Polinton-like virus (PLV) major capsid proteins (9), and single-cell amplified genomes of Chrysophyceae Clade H, encode endogenous viral elements from the *Lavidaviridae* family, which share evolutionary origin and architecture to PLVs, and the *Nucleocytoviricota* phylum (10). Many members of the *Nucleocytoviricota* phylum are referred to as giant viruses (11). Genetic marker surveys of protist and giant virus distributions from the Tara Oceans global dataset also indicate that chrysophytes commonly co-occur with the giant virus family *Mimiviridae* (12). Single-cell RNA-seq studies subsequently provided additional evidence that chrysophytes are native hosts of diverse viruses in the order *Imitervirales* (13). However, chrysophyte-infecting giant virus isolates have not been previously described. In this study, we detail the isolation and initial characterization of a giant virus that infects a Clade H chrysophyte, Chrysophyceae Clade H virus SA1 (ChrysoHV).

Giant viruses encode extensive genomic repertoires that include numerous homologs of cellular genes (14). These cellular homologs may augment or compensate for host functions compromised during infection (15–17) or interfere with host defenses (18). Among the common viral homologs of cellular genes are microbial rhodopsins (19–21), which are photoreceptive proteins that may aid in the generation of ion gradients or intracellular signaling (22, 23). Viral rhodopsin homologs have been experimentally confirmed to convert light energy into electrochemical potential across multiple clades: viral rhodopsin groups I and II (15, 24, 25), channelrhodopsin (16), and heliorhodopsin (18). Furthermore, these viral rhodopsin homologs are globally distributed across aquatic habitats, with their relative abundance decreasing over depth with the attenuation of sunlight (15, 26).

Although giant viruses are diverse and globally abundant (12, 20, 21), a relatively small fraction of their known diversity is represented by cultivated isolates (27). Likewise, giant virus rhodopsin homologs are widespread and potentially functional, but only a few rhodopsin-encoding giant viruses have been isolated and remain in culture with their native host (17, 18, 28). Cultivated isolates are important for establishing virus-host linkages, and furthermore, can reveal novel gene lineages that may go undetected in environmental datasets and clarify the evolutionary history of highly divergent sequences (17). Through the isolation and characterization of ChrysoHV, we identify a previously undescribed lineage of viral rhodopsin that appears to derive from bacterial proteorhodopsin.

## Results & Discussion

### Host range, infection dynamics, and virion structure

We tested the host range of ChrysoHV against seven protists, which were isolated from the same open-ocean site as the virus 100 km north of the island of O‘ahu (Station ALOHA). The protists used in the host range test spanned three taxonomic classes, including three Chrysophyceae, three Dictyochophyceae, and one Bolidophyceae representative. The representative of chrysophyte Clade H lysed when challenged with ChrysoHV (Fig. 1A) and viral particle count increased (Fig. 1B). The apparent burst size, *i.e*., estimated number of virions produced per cell lost, was 24 ± 2 (mean ± SD). No lysis was observed for the other six isolates (Fig. 1B, Fig. S1).

**Fig. 1.**
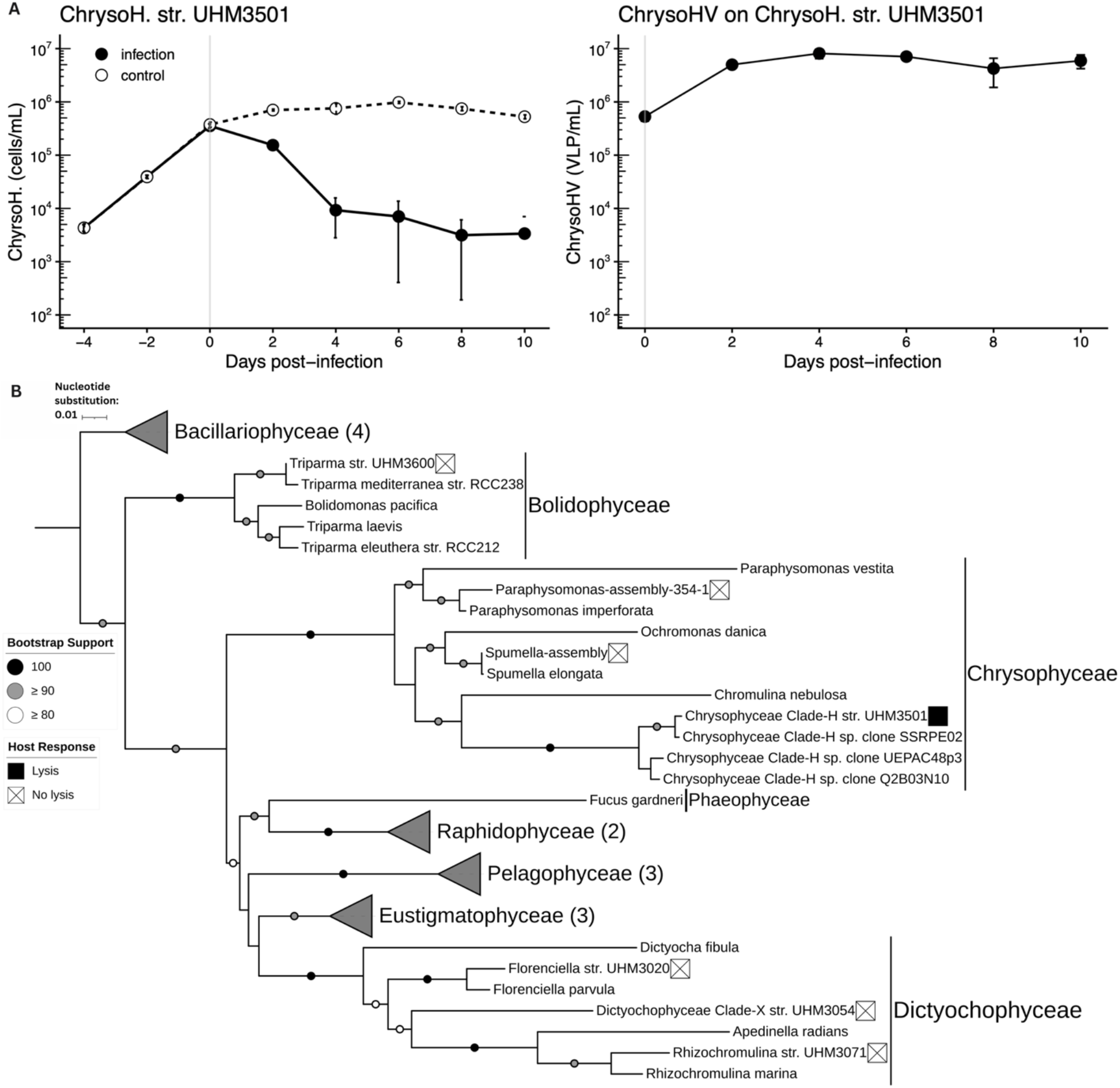
ChrysoHV infection dynamics and host range tests. (A) The left panel shows a time series of Chrysophyceae Clade H sp. UHM3501 abundance (cells/mL) in response to the addition of raw ChrysoHV lysate. The control treatment is shown in the black dashed line with white-fill points, and the infection treatment shown in the solid black line with black-fill points. The right panel shows the abundance of ChrysoHV virus-like particles (VLP/mL) during the same time series. Days pre- and post-infection are displayed on the x-axis. Error bars represent the standard deviation of the triplicate counts. (B) Maximum Likelihood (ML) 18S rRNA phylogeny (GTR+I+R model) of the Ochrophyta phylum with isolates challenged in bold. The white or black boxes by the bolded isolate names indicate whether the protist culture did not lyse (white with “X”) or lysed and produced virions (black) in response to the ChrysoHV challenge. The node symbols indicate the bootstrap values equal to or greater than 80% (white), 90% (gray) or 100% (black). There is no node symbol for branches with less than 80% support.

The ChrysoHV virions displayed an icosahedral capsid (290 ± 40 nm diameter [mean ± SD]) associated with a loose membrane (720 ± 120 nm), and a long tail (1,200 ± 240 nm length, 20 ± 2 width) terminating in tail fibers (200 ± 60 nm length) (Fig. 2A-C; Fig. S2). The capsid structure of ChrysoHV was partially obscured by a sac-like membranous envelope, approximately 710 nm in diameter, always found in association with the negative-stained capsid (Fig. 2A–C, Fig. S3). ChrysoHV’s capsid morphology bears some resemblance to icosahedral capsids of known *Mimiviridae* isolates, and its 290 nm capsid diameter falls within the established range of mimivirus capsid diameters (280-450 nm) (26) (Fig. S2). Transmission electron microscopy (TEM) imaging did not conclusively show whether the membranous envelope structure completely enclosed the capsid or was attached to the capsid vertex. While *Nucleocytoviricota* representatives, notably poxviruses, have lipid envelopes (29), these envelopes are compact and encase the virion core. Marseilleviruses can form large vesicles of viruses packaged inside host-derived membranes, and the resulting membrane-bound particles likely stimulate phagocytosis (30). This strategy of increasing virion volume through an associated membranous structure to enter hosts via phagocytosis may be at play in ChrysoHV infection dynamics. Loosely attached virus envelope structures, referred to as saccule-bearing virus-like particles (VLPs), have been imaged from environmental samples of temperate eutrophic lakes (31). Therefore, although the sac-like envelope is rare among cultivated viruses, it may be geographically widespread.

**Fig. 2.**
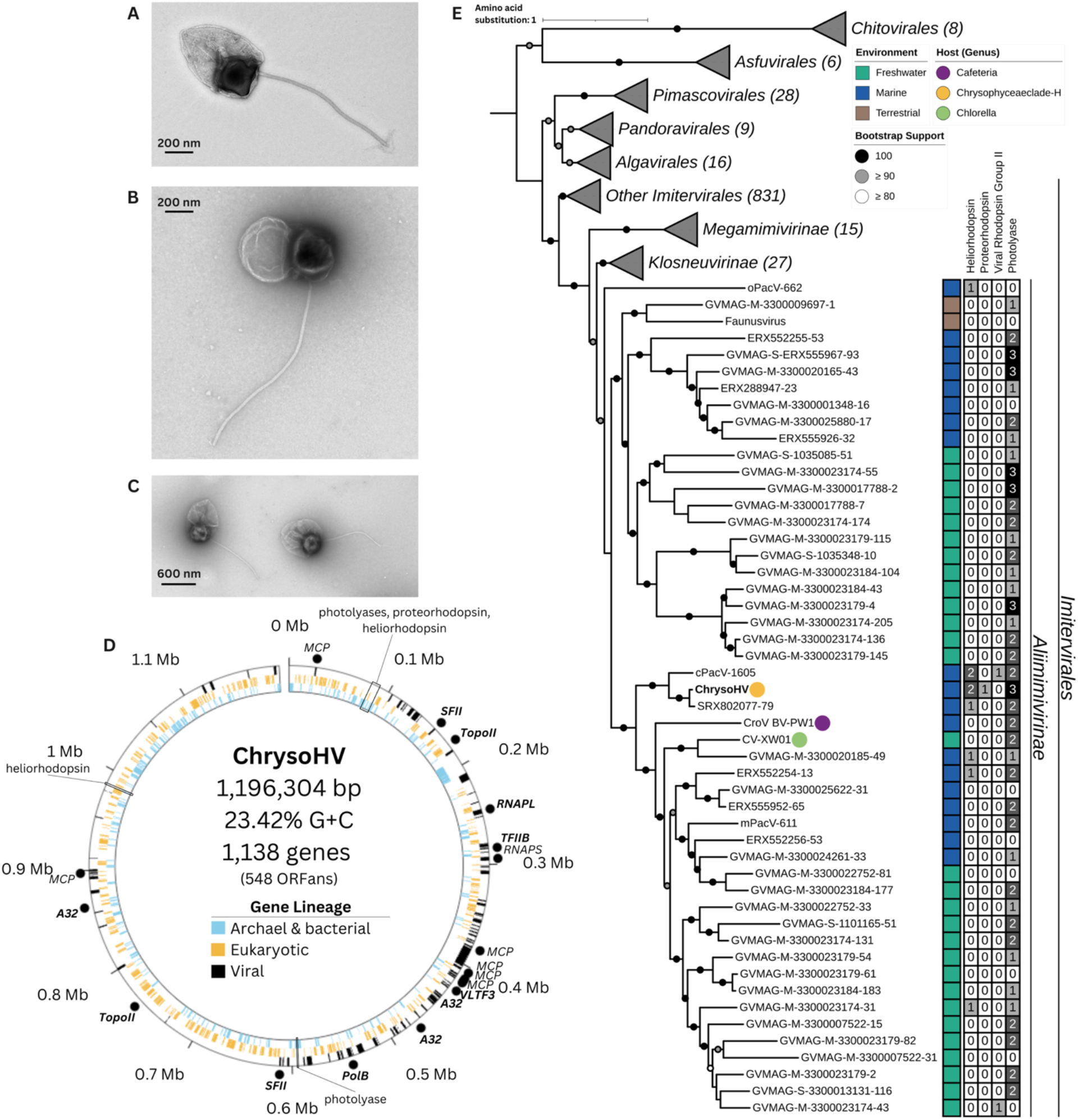
ChrysoHV ultrastructure, genome, and phylogenetic analysis. (A) Transmission electron microscopy (TEM) micrograph of ChrysoHV propagated on UHM3501. (B, C) TEM micrograph of negatively stained ChrysoHV propagated on UHM3500. (D) Genome plot of ChrysoHV with track placement and color showing gene lineage annotated according to domain, inner to outer tracks showing: archaeal and bacterial (blue), eukaryotic (yellow), and viral (black) lineages. Phylogenetically informative giant virus marker genes are represented as black dots on the outside track and show gene name. The genes used in the *Nucleocytoviricota* phylogenetic analysis are bolded. (E) Maximum Likelihood (ML) phylogeny of the *Nucleocytoviricota* phylum inferred from a concatenated alignment of amino acid sequences of marker genes (SFII, RNAPL, PolB, TFIIB, TopoII, A32, VLTF3) using model LG+I+F+G4 with ultrafast bootstrap supports (-bb 1000) shown by node symbol color– white (≥ 80%), gray (≥ 90%), and black (100%) with no node symbol for supports < 80%. The tree is rooted with *Chitovirales* as the outgroup, and the scale bar represents one amino acid substitution per site. The frequency of microbial rhodopsins and photolyases present in *Aliimimivirinae* representatives are annotated in the grayscale matrix, and the environment the representative was collected from is shown in jade (freshwater), blue (marine), and brown (terrestrial).

The micron-length tail structure terminating in tail fibers was observed on most, but not every ChrysoHV virion (Fig. S2), possibly due to the fragility of the tail, which was only 20 nm in width, or the presence of tailed versus un-tailed morphotypes. The amoeba-infecting Naiavirus (32) and Tupanvirus (33), also have long tails that can extend the total virion length beyond 1 µm, and the Naiavirus viral core and tail are completely enclosed by an envelope (32). These other tailed giant viruses enter their host through phagocytosis (32, 33), but the entry mechanism for ChrysoHV has not been determined. Marine giant viruses have been observed with tails extending greater than one micron (34-36), but these virions do not have an associated loose membranous envelope like that of ChrysoHV.

### Genomic characterization

The ChrysoHV genome was assembled as a putatively linear DNA sequence of 1.19 Mbp without terminal inverted repeats, encoding 1,138 predicted genes with a G+C content of 23.42% and 91% estimated completeness (Fig. 2B). ChrysoHV is the third cultivated isolate in the *Aliimimivirinae* subfamily (37) of *Mimiviridae* (Fig. 2E); other known isolates are CroV BV-PW1 (38), infecting a heterotrophic nanoflagellate, and CV-XW01 (39), infecting a green alga. ChrysoHV encodes genes with diverse putative functions, including the ribosomal proteins eL40 (XVM28457.1) and S27a (XVM29017.1), making it the second isolated eukaryote-infecting virus known to encode ribosomal proteins (17), and the first to encode an S27a-like protein (Table S1). ChrysoHV also encodes electron transport proteins such as cytochrome b5, widespread in prokaryotes, eukaryotes, and giant viruses (Table S1) (40), along with succinate dehydrogenase flavoprotein and iron-sulfur subunits (Table S1). While originally thought to be constrained to cellular life, these genes related to the transport of electrons generated from the central carbon metabolism have been detected in metagenome-resolved genomes of giant viruses across aquatic habitats (14, 20, 21).

The ChrysoHV virus encodes seven ABC transporters including one specific to phosphate transport (XVM28869.1) and two AmtB ammonium transporters. Of these ammonium transporters, one (XVM28992.1) has 55% amino acid sequence identity (aaID) to *Prochlorococcus* and *Synechococcus* homologs (Table S1, Fig. S4A). Likewise, ChrysoHV encodes three porins with 44-66% aaID to *Prochlorococcus* porins (XVM28452.1, XVM28569.1, XVM28570.1) (Fig. S4B, C), two urea ABC transporter substrate-binding proteins with 81-90% aaID to a *Prochlorococcus* homolog (XVM29005.1, XVM29006.1) (Fig. S4D), a semiSWEET sugar transporter with 44% aaID to a *Prochlorococcus* homolog (XVM28124.1) (Fig. S4E), a sulfotransferase family protein with 55% aaID to a *Prochlorococcus* homolog (XVM28964.1) (Fig. S4F), and a N-acetylmuramoyl-L-alanine amidase for cell wall remodeling with 39% aaID to a *Prochlorococcus* homolog (XVM29061.1) (Table S1, Fig. S4G). In two of the seven instances in which a ChrysoHV gene had a *Prochlorococcus* homolog, phylogenies of homologous sequences from bacteria, eukaryotes, and viruses showed that the closest relatives derived from a eukaryote (semiSWEET sugar transporter) or a bacterium (N-acetylmuramoyl-L-alanine amidase). However, in most cases (5 of 7), the *Prochlorococcus* homolog was the closest relative to the ChrysoHV gene. This suggests extensive horizontal gene transfer from cyanobacterial prey of the chrysophyte to a giant virus that infects it. Furthermore, phylogenetic placement of two of the porin genes (XVM28569.1, XVM28570.1) and the two urea substrate-binding protein genes (XVM29005.1, XVM29006.1), and their adjacent positions in the ChrysoHV genome (Table S1, Fig. S4), suggest there were gene duplication events after lateral gene transfer.

The ChrysoHV genome also includes a region of putatively light-sensitive proteins in the 85–101 kbp gene neighborhood encompassing a proteorhodopsin-like protein (XVM28022.1), surrounded by two photolyases (XVM28016.1, XVM28025.1), and followed by a heliorhodopsin (XVM28027.1) (Fig. 2D, Fig. 3C, Table S1). The positioning and shared orientation of the photolyases and heliorhodopsin around the proteorhodopsin-like protein implies that they may be part of the same operon. In the bacterium *Trichococcus flocculiformis*, heliorhodopsin and photolyase genes share an operon. In that system, heliorhodopsin binding to photolyase was found to increase photolyase DNA repair activity under green light (41). It is unknown whether the colocalized rhodopsins and photolyases play a similar role in ChrysoHV.

**Fig. 3.**
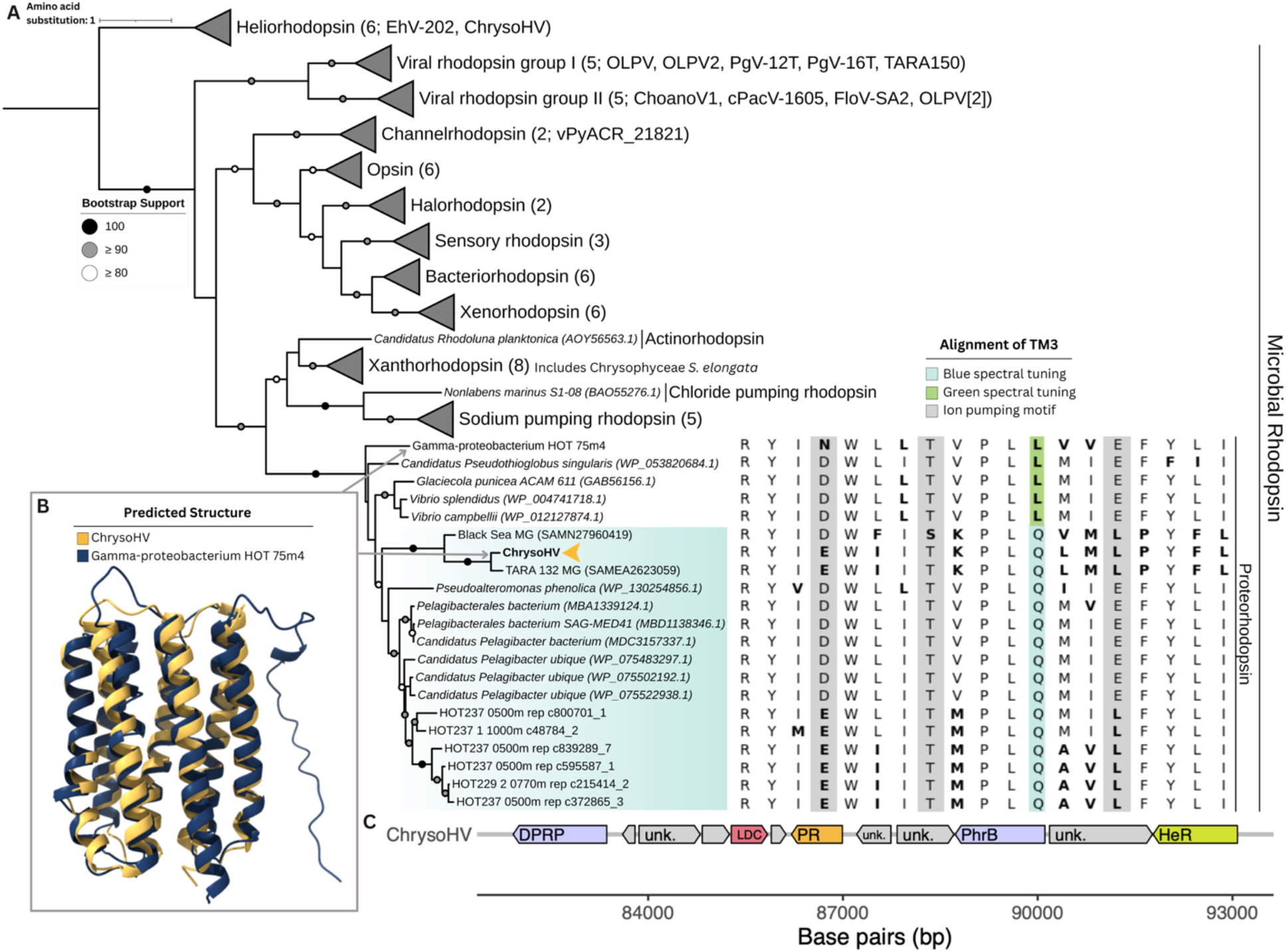
Rhodopsin gene phylogeny, predicted structure, and gene neighborhood. (A) Microbial rhodopsin ML phylogeny with heliorhodopsin as the outgroup, built with model LG+F+R10. Select representatives of viral rhodopsin homologs and putative chrysophyte rhodopsins are referred to in the parenthesis of their corresponding collapsed clade. The proteorhodopsin (PR) clade is annotated with the alignment of transmembrane helix 3 (TM3) outlining the ion-pumping motif (gray) and the spectral tuning amino acid (green or blue). Residues that differ from the most common residue at that locus are in bold (B). The predicted structure of ChrysoHV’s proteorhodopsin-like protein’s conserved domain (yellow) and the HOT 75m4 Gamma-proteobacterium proteorhodopsin (blue). (C) Gene neighborhood of photoreceptive proteins in ChrysoHV, showing the proteorhodopsin (locus tag: ChrysoHV_0101; GenBank ID: XVM28022.1) next to the enzyme lysine decarboxylase (locus tag: ChrysoHV_0099, GenBank ID: XVM28020.1), between two photolyases (locus tag: ChrysoHV_0095, ChyrsoHV_0104; GenBank ID: XVM28016.1, XVM28025.1), and one heliorhodopsin (locus tag: ChrysoHV_0106, GenBank ID: XVM28027.1).

### Sequence characterization of a distinct lineage of proteorhodopsin

In the marine environment, diverse microorganisms from proteobacteria to microalgae use rhodopsins to supplement their energetic needs (42, 43). SAR11 bacteria, of which *Pelagibacter* is an abundant subclade, account for 25% of all plankton (44), and are a likely prey source for mixotrophic algae such as Clade H chrysophytes. Phylogenetic analysis of ChrysoHV’s proteorhodopsin-like protein placed it within the proteorhodopsin clade (Fig. 3A). The sequence alignment of transmembrane domain helix-III (TM3) indicates that the ChrysoHV rhodopsin is spectrally tuned to blue light (glutamine, Q; Fig. 3A), which is the dominant spectral tuning residue in proteorhodopsins from Station ALOHA (42), the isolation site of ChrysoHV. The two most closely related rhodopsin sequences were recovered from metagenomes from the Black Sea at 50 m depth (SAMN27960419) and the North Pacific at 5 m depth in the 0.22–3 μm size fraction (SAMEA2623059). These rhodopsins may also absorb blue light (Fig. 3A) and share a poly-N (asparagine) tract at the start of the rhodopsin coding sequence.

ChrysoHV does not encode the genes needed to synthesize β-carotene, a precursor to retinal, the light-absorbing chromophore in proteorhodopsin. Furthermore, a predicted salt bridge between residues K102 and E228 in the ChrysoHV proteorhodopsin most likely would inhibit retinal binding. This loss of retinal binding capacity may reflect a change in protein function and could explain the high rate of sequence divergence of ChrysoHV’s proteorhodopsin compared to the rest of the proteorhodopsin clade (Fig. 3A). Understanding how the rhodopsin may function without retinal binding could be important to interpreting the function of viral rhodopsins detected below the photic zone (45, 46). Further investigation is needed to confirm whether the ChrysoHV proteorhodopsin can confer other physiological benefits in the absence of light absorption, similar to the *Psychroflexus torquis* proteorhodopsin ptqPR, which acts as a phospholipid scramblase to preserve membrane potential, fluidity, and integrity (47).

The ChrysoHV proteorhodopsin also lacks the conserved DTE proton pumping motif found in the third transmembrane domain of most proteorhodopsins, and instead, possesses an ETL motif (Fig. 3A), which has been detected in proteorhodopsins from deep water at Station ALOHA (Fig. 3A) (42). A comparison of the predicted structure of the conserved domain of the ChrysoHV rhodopsin against existing functional characterizations revealed that it was most similar to the uncultured gamma-proteobacterium clone HOT 75m4 rhodopsin (Fig. 3B), although the relatively low amino acid identity (35.9% aaID) limits functional inference. The latter rhodopsin was confirmed to pump protons (H^+^) (48), and, like ChrysoHV, was collected from Station ALOHA. These results suggest that ChrysoHV may have acquired the proteorhodopsin from a marine bacterium, which, like *Prochlorococcus*, may be prey for chrysophyte Clade H.

### Comparison to photoreceptive proteins across Nucleocytoviricota

In the *Aliimimivirinae* subfamily, only two other representatives, cPacV-1605 and GVMAG-M-3300023174-43, encode rhodopsins, both belonging to viral rhodopsin group II (49) (Fig. 2A). However, 40 other *Aliimimivirinae* representatives encode at least one photolyase homolog (Fig. 2E), either the (6-4) photolyase or DASH-type cryptochrome. Only ChrysoHV and cPacV-1605 encode multiple rhodopsin and photolyase types, including two heliorhodopsins (16, 49) (Fig. 2E). However, cPacV-1605 encodes a viral rhodopsin (group II) (49), and ChrysoHV encodes a proteorhodopsin (Fig. 2E, 3A). Furthermore, instead of being nested within the putatively photosensitive gene neighborhood, the position of the viral rhodopsin in cPacV-1605 (locus tag: CPAV1605_207, GenBank ID: VVU94485.1) is further away from the photolyases (locus tag: CPAV1605_133, CPAV1605_136; GenBank ID: VVU94411.1, VVU94414.1) and heliorhodopsins (locus tag: CPAV1605_533, CPAV1605_1309; GenBank ID: VVU95557.1, VVU94808.1) (49). ChrysoHV is the only cultivated giant virus known so far to encode a proteorhodopsin-lineage rhodopsin along with two heliorhodopsins (Fig 2E, Fig. 3A).

### Global ocean distribution of ChrysoHV phylotypes and proteorhodopsins

Analysis of the distribution of ChrysoHV phylotypes was determined by identifying similar DNA polymerase family B amino acid sequences (≥ 90% PolB aaID), which are a highly conserved *Nucleocytoviricota* marker genes (11), from within the *Tara Oceans* dataset (12). This analysis revealed the prevalence of ChrysoHV phylotypes in subtropical surface ocean sites in the Pacific and Indian Ocean as well as the Mediterranean Sea (Fig. 4A). Notably, at station 132 north of the Hawaiian archipelago, ChrysoHV phylotypes were detected in both the mixed layer and mesopelagic zone, suggesting connectivity between surface and deep ocean virus populations, a connection previously observed in the tropical North Pacific (26). ChrysoHV proteorhodopsin-like proteins (≥ 80% aaID) were detected at fewer stations than ChrysoHV-like PolBs, but their relative abundance was also highest in the subtropical surface ocean at Station 132 (Fig. 4B) suggesting that some of these sequences derive from ChrysoHV-like viruses.

**Fig. 4.**
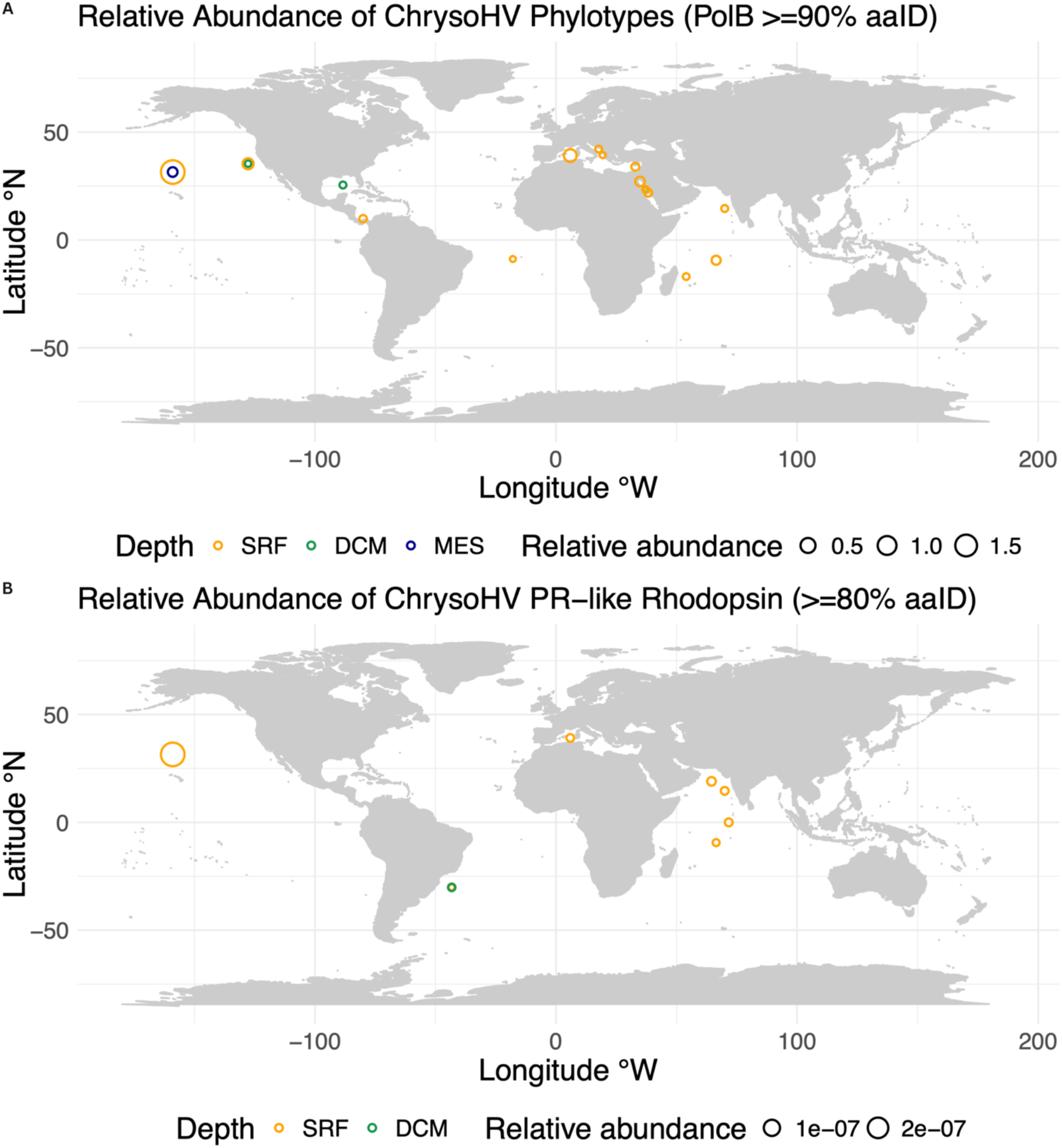
Distribution and relative abundance of ChrysoHV PolB (DNA polymerase family B) and PR (proteorhodopsin). (A) Global distribution of phylotypes with ≥ 90% PolB aaID to ChrysoHV from the *Tara* Oceans dataset. Size of the circle indicates the relative abundance of the summed phylotypes at each given sample site to total *Nucleocytoviricota* PolB sequences per site. (B) Global distribution of homologous PR with ≥ 80% aaID to the ChrysoHV proteorhodopsins with size of the circle indicating relative abundance normalized to total metagenomic reads per site. Sample depth is shown through color: surface (SRF: 3-7 m, orange), deep chlorophyll maximum (DCM: 7-200 m, green), and mesopelagic (MES: 200-1,000 m, purple).

## Conclusion

The phytoplankton-infecting virus ChrysoHV is unusual in its morphology, with a long flexible tail and sac-like membrane that expands the documented phenotypic diversity of the virosphere. It is also unusual in its genetic composition. As the only cultivated virus isolate known to encode multiple lineages of rhodopsins that fall within clades from proteorhodopsin to heliorhodopsin, and only the second isolate known to encode ribosomal proteins, ChrysoHV offers an opportunity for experimental work on what may be novel mechanisms for manipulating host physiology during giant virus infections. The discovery of multiple bacteria- and cyanobacteria-derived genes in ChrysoHV supports the proposition that protists are a locus of direct or indirect lateral gene transfer between entities that would not otherwise interact, namely, a eukaryotic virus and a bacterium (50). The internalization by phagotrophic protists of genomic DNA from both an infecting virus and ingested bacterial prey suggests that lateral transfer could occur directly between bacterium and virus, but it may also be mediated indirectly by exchanges from bacterial prey to protist, then protist to virus. This experimentally accessible virus-host system may prove useful for future investigations into how the virocell environment of predatory protists influences gene acquisition in giant viruses.

## Materials and Methods

### Isolation and cultivation

The five mixotrophic protists used in this study include two chrysophytes (UHM3500, UHM3501), one bolidophyte (UHM3600), and three dictyochophytes (UHM3020, UHM3053, UHM3071), all of which were isolated from the epipelagic zone (5–100m) in the North Pacific Subtropical Gyre at Station ALOHA (22° 45’ N, 158° W) through serial dilutions of whole seawater samples enriched with *Prochlorococcus* (7). The mixotrophs were cultivated at 24°C at approximately 200–220 µmol photons m^-2^ s^-1^ irradiance to mimic the high light conditions at Station ALOHA (51). The unialgal cultures were grown on K medium without dissolved, inorganic nitrogen sources (NaNO_3_ and NH_4_Cl) (52). Cultures were instead amended with cyanobacteria prey (*Prochlorococcus* MIT9301) for their nitrogen source (7, 53). In addition, two heterotrophic chrysophytes (UHM3530 and UHM3560) were used in this study. These strains were isolated from seawater collected from the mixed layer (5–50 m) at Station ALOHA in 2022 and 2024 through serial dilutions of seawater samples. The heterotrophs were cultivated on filtered seawater amended with sterile quinoa grains to stimulate growth of ambient bacteria as their food source.

The Chrysophyceae Clade H virus SA1 (ChrysoHV) was isolated by challenging the chrysophyte isolate UHM3500 with unfiltered seawater collected from 25 m depth at Station ALOHA on 04 May 2019. The lytic agent detected in the challenge was rendered clonal by serial dilutions to extinction (54). This is the first chrysophyte-infecting virus isolated from Station ALOHA, thus the strain designation SA1. In addition to the host used for isolation, the virus was found to infect a related chrysophyte strain UHM3501 (Fig. S1). Both strains are members of Chrysophyceae Clade H with nearly identical 18S rRNA sequences and similar grazing kinetics on *Prochlorococcus* (7). After discovering a heterotrophic protist contaminant in UHM3500, the virus was propagated exclusively on UHM3501. The virus was maintained by infecting the host alga (25 mL) in exponential growth phase with archived lysate from a previous infection (0.5 mL) approximately monthly. At three to four days post-infection (DPI), the infected culture was stored at 4°C to serve as lysate for the next round of infection.

### Host range tests and flow cytometry

We challenged isolates of Ochrophyta protists, both mixotrophs and heterotrophs, with raw ChrysoHV lysate, then tracked the population dynamics of the protists and viruses over time using flow cytometry. The phago-mixotrophic protists were tracked every two days up to ten DPI, and the heterotrophic protists were tracked every day up to five DPI. The experimental design included triplicate, 25 mL protist cultures for both negative controls and infection treatments. To prepare the infection treatment, 500 μL of unfiltered lysate (3-6 × 10⁵ virus-like particles mL⁻¹ final concentration) was added to exponential or early stationary phase protist cultures. The samples were fixed with ice-cold glutaraldehyde at 0.5% final concentration, incubated for 15 min at 4°C, and then flash frozen in liquid nitrogen (7, 53).

The samples were stored at -80°C up to four days, then thawed at room temperature and processed within the same day on a CytoFlex-S (Beckman Coulter Life Sciences, Indianapolis, IN, USA) in the Biological Electron Microscope Facility (BEMF) at the University of Hawai‘i at Mānoa. The samples were split into two fractions, with 200 μL allocated towards quantifying cell abundances and loaded directly onto a 96-well plate, and 5 μL allocated toward quantifying virus abundances. The latter fractions were diluted into 195 μL solutions of TE (pH 8.0) with 2.5X SYBR Green I nucleic acid stain (Invitrogen, Thermo Fisher Scientific, Waltham, MA, USA) and incubated at 80°C for 20 min in darkness (55), then loaded onto a 96-well plate. The cell counts were run on auto-record for one minute at a flow rate of 20 μL min^-1^ for the algae and 30 μL min^-1^ for the heterotrophs. The mixotroph populations were distinguished through chlorophyll fluorescence and 488 nm, 90° side scatter (SSC), whereas the heterotrophs were detected using SYBR Green fluorescence and 488 nm, 90° SSC (56). The VLP population was identified through SYBR Green fluorescence and 405 nm, 90° violet side scatter (VSSC), enhancing marine virus detection (57). The samples were run on auto-record for one minute at a flow rate of 10 μL min^-1^. Both algae and virus populations were delineated through a custom Python script using the following packages: fcsparser (v0.2.8), pandas (v2.2.2), matplotlib (v3.7.1), while the FlowJo (v.10.10.0) software was used to delineate the heterotrophic protist population. Apparent burst size was determined by dividing the maximum concentration of VLPs post-infection by the inferred number of lysed cells, calculated as the difference between the cell concentration on the day of infection minus the minimum concentration post-infection.

### Electron microscopy

TEM micrographs of viral particles were prepared from raw, unfixed lysate and negatively stained with 2% uranyl acetate for 45 sec on 200-mesh copper grids with carbon-stabilized formvar support that had been rendered hydrophilic by glow discharge (58). Images were captured with a Hitachi HT7700 transmission electron microscope (Tokyo, Japan) at the Biological Electron Microscope Facility (BEMF) at the University of Hawai‘i at Mānoa. Particle dimensions were measured with ImageJ from Fiji (v2.16.0/1.54p) (59), including: capsid diameter (n = 38), envelope (n = 34), tail (n = 17), and tail fibers (n = 8).

### DNA extraction, sequencing, assembly, and annotation

To prepare sufficient viral biomass for DNA sequencing, DNA was extracted from 610 mL of lysate (6.4 × 10^6^ VLP/mL), after allowing the infection of chrysophyte Clade H sp. UHM3501 to clear four DPI. The unfiltered lysate was concentrated and pooled through serial centrifugations (6X) in a 100 kDA Centricon Plus centrifugal filter (MilliporeSigma, Burlington, MA, USA) at 3,500 × *g* at 20°C for 15 minutes. Approximately 200 ng of DNA was extracted from the concentrate using the ZymoBiomics Quick-DNA High Molecular Weight MagBead Kit (Zymo Research, Irvine, CA, USA), flash frozen in liquid nitrogen, stored at -20°C, then shipped on dry ice to the US Department of Energy Joint Genome Institute (JGI). There, the DNA was sequenced using PacBio Multiplexed 6–10 kb Low Input parameters.

A PacBio Low Input (DNA) library was constructed and sequenced using the PacBio SEQUELIIe platform (Pacific Biosciences, Menlo Park, CA USA), which generated 3,267,906 reads totaling 31,033,080,767 bp. Reads were deduplicated using pbmarkdup (v1.0.3) (60). BBDuk (v38.99) (61) was used to remove reads that contained PacBio control sequences and PCR adapters, and IceCreamFinder (v38.99) (62) was used to remove reads without SMRTbell sequences. The final filtered fastq contained 3,257,268 reads totaling 30,921,351,264 bp with mean read length of 9,493 bp.

The filtered reads (31 Gb) were assembled using Flye (v2.9) (63) with the “--meta --pacbio-hifi” argument with a minimum read overlap of 7,000 bp. From this initial assembly, viral contigs were identified and grouped through a low-complexity binning workflow, utilizing CONCOCT (v1.1.0) (64), MaxBin (v2.2.7) (65), MetaBAT2 (v2.15-5) (66), and SemiBin2 (v1.5.1) (67). For each binning strategy, the viral bin was identified by assessing the presence of giant virus marker genes (11) with GVClass (v1.0) (68), and the MetaBAT2-generated viral bin was used in downstream analysis. Subsequently, circular consensus sequencing reads were recruited against these identified viral contigs, and a 100 Mb subsample of these reads was assembled using Flye without the “--meta” argument (v2.9) (63). This iterative assembly resulted in one high-quality viral contig, which was polished using two rounds of Racon (v1.6.0) (69) to ensure accuracy.

Gene calling and estimated completeness was performed with GVClass (v1.0) (65), and the 1.196 Mb viral genome had an estimated 93.77% coding density. Gene annotation was performed with KofamScan (v1.3.0) (70) and EggNOG-mapper (v2.1.12) against the EggNOG 6.0 database (71) with annotations of genes of interest confirmed with BLASTp (72) against the NCBI ClusteredNR database using the protein-protein BLAST algorithm. tRNAs were annotated with tRNAscan-SE (v2.0.12) (73).

### Phylogenetic analysis

Phylogenetic analysis of ChrysoHV with IQ-TREE (v2.2.6) (74) was based on a concatenated alignment of seven core proteins: SFII, RNAPL, PolB, TFIIB, TopoII, A32, and VLTF3 (11). Select *Nucleocytoviricota* sequences were retrieved from the GVMAGs (v1) (21) and GenBank Assembly (75) databases, and quality filtered to contain at least four out of seven core proteins determined through hmmsearch (v3.3.2, E-value ≤ 1 × 10^-10^) (76). Additionally, the query sequences were filtered to remove genomes with more than 10 copies of any core protein and more than four core proteins with multiple copies. The remaining 990 query sequences were aligned with MAFFT (v7.520) (77) and trimmed with trimAl (-gt 0.1; v1.4.1) (78), then concatenated with a custom python script (79). The trimmed, concatenated alignment was used for phylogenetic analysis using the mixture model LG+I+F+G4 and ultrafast bootstrap supports (option: -bb 1000) (v2.2.6) (74). The tree was annotated with iTOL (v6) (80).

Gene phylogeny of microbial rhodopsins with a heliorhodopsin outgroup were built with IQ-TREE (v2.2.6) (74) using model LG+F+R10 and ultrafast bootstrap supports (option: -bb 1000) and annotated with iTOL (v6) (80). Representatives of rhodopsin homologs from cultivated virus isolates include: EhV-202 (heliorhodopsin) (17), PgV-12T and -16T (viral rhodopsin I) (27), and FloV-SA2 (viral rhodopsin II) (16). Additionally, the gene phylogeny includes synthetic construct: vPyACR_21921 (channelrhodopsin) (15), and rhodopsin homologs from single amplified genomes (SAGs) and metagenome assembled genomes (MAGS): ChoanoV1 (viral group II) (14), cPacV-1605 (viral group II) (49), and OLPV 1 and 2 (viral groups I & II) (81). Query rhodopsin sequences were retrieved from NCBI and the supplementary materials of Olson *et al*. (2018), which included proteorhodopsin sequences recovered from metagenomic sequences at Station ALOHA (42). The alignment was built with MAFFT (L-INS-i; v7.520) (77) and trimmed with trimAl (-gt 0.1; v1.4.1) (78).

The phylogeny of near-complete 18S rRNA genes from select algal isolates were built with IQ-TREE (v2.2.6) (74) using GTR+I+G model with ultrafast bootstrap support (option: -bb 1000) and annotated with iTOL (v6) (80). The nucleotide sequences were retrieved from NCBI, aligned with MAFFT (L-INS-i; v7.520) (77) and trimmed with trimAl (-gt 0.1; v1.4.1) (78).

### Gene distribution in the global ocean

Comparisons to *Nucleocytoviricota* PolB amino acid sequences from the Tara Oceans dataset (12) were performed using BLASTp (v2.13.0, E-value ≤ 1 × 10^-120^) (72), and sequences with 90% or greater aaID were plotted using a custom R script with the following packages: maps (v3.4.2), mapdata (v2.3.1), and ggplot2 (v.3.5.1). The relative abundance of the PolB genes were normalized to the total number of *Nucleocytoviricota* PolBs (12). Likewise, ChrysoHV proteorhodopsin-like protein sequences from the Tara Oceans dataset with a minimum of 80% aaID were identified through Ocean Gene Atlas (82, 83). The relative abundance of ChrysoHV proteorhodopsin-like protein sequences were normalized to total metagenomic reads in the given sample (82, 83).

### Protein structure predictions

The conserved domain of the ChrysoHV rhodopsin structure was predicted using ColabFold (v1.5.5) (84) and compared against the AlphaFold Protein Structure Databases–SwissProt (85) and CATH (v4.4) (86) using Foldseek (87). The conserved domain of the ChrysoHV rhodopsin and HOT 75m4 Gamma-proteobacterium proteorhodopsin structures were rendered in ChimeraX (v1.8.dev202403010247) (88).

## Acknowledgments

We thank the personnel of the Hawai‘i Ocean Time-series program (NSF award 12-60164) and R/V *Kilo Moana* for assistance with seawater collection. The Biological Electron Microscope Facility (BEMF) at the University of Hawai‘i at Mānoa was instrumental in this research and we acknowledge the facility staff, Tina Weatherby and Dr. Orion Rivers, for their technical assistance. We are indebted to Kelsey McBeain for maintenance of the UHM library culture collection, and US DOE Joint Genome Institute Scientific Project Manager, Kerrie Barry, for overseeing sequencing of ChrysoHV. We are grateful for thoughtful feedback provided by Dr. Karen Selph (Department of Oceanography, University of Hawai‘i at Mānoa) and Dr. Alison Sherwood (School of Life Sciences, University of Hawai‘i at Mānoa) on manuscript preparation, and for the insights into rhodopsin structure and rhodopsin sequences (SAMN27960419 and SAMEA2623059) provided by Dr. Oded Béjà and Dr. Andrey Rozenberg (Faculty of Biology, Technion-Israel Institute of Technology).

## Author Contributions

GFS, KFE, and PB developed, designed, and acquired funding support for the project. CRS isolated the virus and algae strains. CW and QL assisted with the collection of virus host-range data. FS provided sequencing support and supervised the bioinformatics analyses. KL assisted with the virus genome assembly. JT assisted with gene phylogeny workflows. CRS and PB imaged the virus. PB wrote the manuscript with input from all authors.

## Data Availability

ChrysoHV sequencing data is available through the NCBI sequence read archive SRR34002084, and the genome assembly has been deposited to NCBI GenBank under accession ID PV764857. The mixotrophic protists used in the study, GenBank IDs: MZ611734.1 (Chrysophyceae Clade H sp. UHM3500), MZ611720.1 (Chrysophyceae Clade H sp. UHM3501), MN615710.1 (*Florenciella* sp. UHM3020), MZ611733.1 (Dictyochophyceae clade X sp. UHM3054), MZ611705.1 (*Rhizochromulina* sp. UHM3071), MZ611730.1 (*Triparma* sp. UHM3600), are available through NCMA. The virus and heterotrophic protists used in this study, GenBank IDs: PV717318.1 (*Paraphysomonas* sp. UHM3560) and PV717319.1 (*Spumella* sp. UHM3530), are available through the Marine Viral Ecology Laboratories at the University of Hawai‘i at Mānoa upon request to KFE and GFS. Additional data and scripts on gene alignments, phylogenies, genome annotation, protein structural predictions, host range challenges, and biogeography are available on Figshare, doi:10.6084/m9.figshare.33099290.

## References

1. Del Campo J, Massana R. 2011. Emerging diversity within chrysophytes, choanoflagellates and bicosoecids based on molecular surveys. Protist 162:435–448.

2. Seeleuthner Y, Mondy S, Lombard V, Carradec Q, Pelletier E, Wessner M, Leconte J, Mangot J-F, Poulain J, Labadie K, Logares R, Sunagawa S, De Berardinis V, Salanoubat M, Dimier C, Kandels-Lewis S, Picheral M, Searson S, Tara Oceans Coordinators, Acinas SG, Boss E, Follows M, Gorsky G, Grimsley N, Karp-Boss L, Krzic U, Not F, Ogata H, Raes J, Reynaud EG, Sardet C, Speich S, Stemmann L, Velayoudon D, Weissenbach J, Pesant S, Poulton N, Stepanauskas R, Bork P, Bowler C, Hingamp P, Sullivan MB, Iudicone D, Massana R, Aury J-M, Henrissat B, Karsenti E, Jaillon O, Sieracki M, De Vargas C, Wincker P. 2018. Single-cell genomics of multiple uncultured stramenopiles reveals underestimated functional diversity across oceans. Nat Commun 9:310.

3. Olefeld JL, Majda S, Albach DC, Marks S, Boenigk J. 2018. Genome size of chrysophytes varies with cell size and nutritional mode. Org Divers Evol 18:163–173.

4. Dai M, Luo Y, Achterberg EP, Browning TJ, Cai Y, Cao Z, Chai F, Chen B, Church MJ, Ci D, Du C, Gao K, Guo X, Hu Z, Kao S, Laws EA, Lee Z, Lin H, Liu Q, Liu X, Luo W, Meng F, Shang S, Shi D, Saito H, Song L, Wan XS, Wang Y, Wang W, Wen Z, Xiu P, Zhang J, Zhang R, Zhou K. 2023. Upper ocean biogeochemistry of the oligotrophic North Pacific Subtropical Gyre: from nutrient sources to carbon export. Reviews of Geophysics 61:e2022RG000800.

5. Frias-Lopez J, Thompson A, Waldbauer J, Chisholm SW. 2009. Use of stable isotope-labelled cells to identify active grazers of picocyanobacteria in ocean surface waters. Environmental Microbiology 11:512–525.

6. Wilken S, Yung CCM, Poirier C, Massana R, Jimenez V, Worden AZ. 2023. Choanoflagellates alongside diverse uncultured predatory protists consume the abundant open-ocean cyanobacterium *Prochlorococcus*. Proc Natl Acad Sci USA 120(27):e2302388120.

7. Li Q, Edwards KF, Schvarcz CR, Steward GF. 2022. Broad phylogenetic and functional diversity among mixotrophic consumers of *Prochlorococcus*. The ISME Journal 16:1557–1569.

8. Flombaum P, Gallegos JL, Gordillo RA, Rincón J, Zabala LL, Jiao N, Karl DM, Li WKW, Lomas MW, Veneziano D, Vera CS, Vrugt JA, Martiny AC. 2013. Present and future global distributions of the marine Cyanobacteria *Prochlorococcus* and *Synechococcus*. Proc Natl Acad Sci USA 110:9824–9829.

9. Bellas C, Hackl T, Plakolb M-S, Koslová A, Fischer MG, Sommaruga R. 2023. Large-scale invasion of unicellular eukaryotic genomes by integrating DNA viruses. Proc Natl Acad Sci USA 120(16):e2300465120

10. Castillo YM, Mangot J, Benites LF, Logares R, Kuronishi M, Ogata H, Jaillon O, Massana R, Sebastián M, Vaqué D. 2019. Assessing the viral content of uncultured picoeukaryotes in the global-ocean by single cell genomics. Molecular Ecology 28:4272–4289.

11. Aylward FO, Moniruzzaman M, Ha AD, Koonin EV. 2021. A phylogenomic framework for charting the diversity and evolution of giant viruses. PLoS Biol 19:e3001430.

12. Endo H, Blanc-Mathieu R, Li Y, Salazar G, Henry N, Labadie K, De Vargas C, Sullivan MB, Bowler C, Wincker P, Karp-Boss L, Sunagawa S, Ogata H. 2020. Biogeography of marine giant viruses reveals their interplay with eukaryotes and ecological functions. Nat Ecol Evol 4:1639–1649.

13. Fromm A, Hevroni G, Vincent F, Schatz D, Martinez-Gutierrez CA, Aylward FO, Vardi A. 2024. Single-cell RNA-seq of the rare virosphere reveals the native hosts of giant viruses in the marine environment. Nat Microbiol 9:1619–1629.

14. Moniruzzaman M, Erazo Garcia MP, Farzad R, Ha AD, Jivaji A, Karki S, Sheyn U, Stanton J, Minch B, Stephens D, Hancks DC, Rodrigues RAL, Abrahão JS, Vardi A, Aylward FO. 2023. Virologs, viral mimicry, and virocell metabolism: the expanding scale of cellular functions encoded in the complex genomes of giant viruses. FEMS Microbiology Reviews 47:fuad053.

15. Needham DM, Yoshizawa S, Hosaka T, Poirier C, Choi CJ, Hehenberger E, Irwin NAT, Wilken S, Yung C-M, Bachy C, Kurihara R, Nakajima Y, Kojima K, Kimura-Someya T, Leonard G, Malmstrom RR, Mende DR, Olson DK, Sudo Y, Sudek S, Richards TA, DeLong EF, Keeling PJ, Santoro AE, Shirouzu M, Iwasaki W, Worden AZ. 2019. A distinct lineage of giant viruses brings a rhodopsin photosystem to unicellular marine predators. Proc Natl Acad Sci USA 116:20574–20583.

16. Rozenberg A, Oppermann J, Wietek J, Fernandez Lahore RG, Sandaa R-A, Bratbak G, Hegemann P, Béjà O. 2020. Lateral gene transfer of anion-conducting channelrhodopsins between green algae and giant viruses. Current Biology 30:4910–4920.e5.

17. Thomy J, Schvarcz CR, McBeain KA, Edwards KF, Steward GF. 2024. Eukaryotic viruses encode the ribosomal protein eL40. npj Viruses 2:51.

18. Hososhima S, Mizutori R, Abe-Yoshizumi R, Rozenberg A, Shigemura S, Pushkarev A, Konno M, Katayama K, Inoue K, Tsunoda SP, Béjà O, Kandori H. 2022. Proton-transporting heliorhodopsins from marine giant viruses. eLife 11:e78416.

19. Yutin N, Koonin EV. 2012. Proteorhodopsin genes in giant viruses. Biol Direct 7:34.

20. Moniruzzaman M, Martinez-Gutierrez CA, Weinheimer AR, Aylward FO. 2020. Dynamic genome evolution and complex virocell metabolism of globally-distributed giant viruses. Nat Commun 11:1710.

21. Schulz F, Roux S, Paez-Espino D, Jungbluth S, Walsh DA, Denef VJ, McMahon KD, Konstantinidis KT, Eloe-Fadrosh EA, Kyrpides NC, Woyke T. 2020. Giant virus diversity and host interactions through global metagenomics. Nature 578:432–436.

22. Pinhassi J, DeLong EF, Béjà O, González JM, Pedrós-Alió C. 2016. Marine bacterial and archaeal ion-pumping rhodopsins: genetic diversity, physiology, and ecology. Microbiol Mol Biol Rev 80:929–954.

23. Rozenberg, A Inoue, K, Kandori, H, Béjà O. Microbial rhodopsins: the last two decades. Annu Rev Microbiol 75:427–447.

24. Bratanov D, Kovalev K, Machtens J-P, Astashkin R, Chizhov I, Soloviov D, Volkov D, Polovinkin V, Zabelskii D, Mager T, Gushchin I, Rokitskaya T, Antonenko Y, Alekseev A, Shevchenko V, Yutin N, Rosselli R, Baeken C, Borshchevskiy V, Bourenkov G, Popov A, Balandin T, Büldt G, Manstein DJ, Rodriguez-Valera F, Fahlke C, Bamberg E, Koonin E, Gordeliy V. 2019. Unique structure and function of viral rhodopsins. Nat Commun 10:4939.

25. Zabelskii D, Alekseev A, Kovalev K, Rankovic V, Balandin T, Soloviov D, Bratanov D, Savelyeva E, Podolyak E, Volkov D, Vaganova S, Astashkin R, Chizhov I, Yutin N, Rulev M, Popov A, Eria-Oliveira A-S, Rokitskaya T, Mager T, Antonenko Y, Rosselli R, Armeev G, Shaitan K, Vivaudou M, Büldt G, Rogachev A, Rodriguez-Valera F, Kirpichnikov M, Moser T, Offenhäusser A, Willbold D, Koonin E, Bamberg E, Gordeliy V. 2020. Viral rhodopsins 1 are an unique family of light-gated cation channels. Nat Commun 11:5707.

26. Sheam MM, Luo E. 2025. Vertical transport and spatiotemporal dynamics of giant viruses in the North Pacific subtropical gyre. The ISME Journal 19:wraf094.

27. Bosmon T, Abergel C, Claverie J-M. 2025. 20 years of research on giant viruses. npj Viruses 3:9.

28. Brussaard CPD, Bratbak G, Baudoux A-C, Ruardij P. 2007. Phaeocystis and its interaction with viruses. Biogeochemistry 83:201–215.

29. Moss B. 2018. Origin of the poxviral membrane: a 50-year-old-riddle. 2018. PLoS Pathog 14:e1007002G.

30. Arantes TS, Rodrigues RA, Dos Santos Silva LK, Oliveira GP, de Souza HL, Khalil JY, de Oliveira DB, Torres AA, da Silva LL, Colson P, Kroon EG, da Fonseca FG, Bonjardim CA, La Scola B, Abrahão JS. 2016. The large marseillevirus explores different entry pathways by forming giant infectious vesicles. J Virol 12:90(11):5246-5255.

31. Billard H, Fuster M, Enault F, Carrias J-F, Fargette L, Carrouée M, Desmares P, Delmont TO, Bigeard E, Tanguy G, Nogaret P, Baudoux A-C, Christaki U, Sime-Ngando T, Colombet J. 2024. Unexpected diversity and ecological significance of uncultivable large virus-like particles in aquatic environments. ISME Commun 5:ycaf098.

32. Rodrigues M, Queiroz V, Arantes T, Limboço H., Neiva B, Arias N, Machado T, Barcelos M, Cortines J, Thiemann OH, Marques RE, Melo-Hanchuk TD, Celis EHL, Araujo JP, Reis E, Alcantara LC, Santos CBC, Jivaji A, Rodrigues RAL, Aylward FO, Abrahão JS. 2025. Naiavirus: an enveloped giant virus with a pleomorphic, flexible tail. Nat Commun 16: 8306.

33. Abrahão J, Silva L, Silva LS, Khalil JYB, Rodrigues R, Arantes T, Assis F, Boratto P, Andrade M, Kroon EG, Ribeiro B, Bergier I, Seligmann H, Ghigo E, Colson P, Levasseur A, Kroemer G, Raoult D, La Scola B. 2018. Tailed giant Tupanvirus possesses the most complete translational apparatus of the known virosphere. Nat Commun 9:749.

34. Bratbak G, Haslund OH, Heldal M, Næss A, Røeggen T. 1992. Giant marine viruses? Mar Ecol Prog Ser 85:201–202.

35. Cochlan WP, Wikner J, Steward GF, Smith DC, Azam, F. 1993. Spatial distribution of viruses, bacteria and chlorophyll a in neritic, oceanic and estuarine environments. Mar Ecol Prog Ser 92:77–87.

36. Gowing M, Riggs B, Garrison D, Gibson A, Jeffries M. 2002. Large viruses in Ross Sea late autumn pack ice habitats. Mar Ecol Prog Ser 241:1–11.

37. Aylward FO, Abrahão JS, Brussaard CPD, Fischer MG, Moniruzzaman M, Ogata H, Suttle CA. 2023. Taxonomic update for giant viruses in the order Imitervirales (phylum *Nucleocytoviricota*). Arch Virol. 168:283

38. Fischer MG, Allen MJ, Wilson WH, Suttle CA. 2010. Giant virus with a remarkable complement of genes infects marine zooplankton. Proc Natl Acad Sci USA 107:19508–19513.

39. Sheng Y, Wu Z, Xu S, Wang Y. 2022. Isolation and identification of a large green alga virus (*Chlorella* Virus XW01) of *mimiviridae* and its virophage (*Chlorella* Virus Virophage SW01) by using unicellular green algal cultures. J Virol 96:e02114–21.

40. Lamb DC, Goldstone JV, Belhaouari DB, Andréani J, Farooqi A, Allen MJ, Kelly SL, La Scola B, Stegeman JJ. 2025. Cytochrome b5 occurrence in giant and other viruses belonging to the phylum Nucleocytoviricota. npj Viruses 3:8.

41. Shim J, Cho S-G, Kim S-H, Chuon K, Meas S, Choi A, Jung K-H. 2022. Heliorhodopsin helps photolyase to enhance the DNA repair capacity. Microbiol Spectr 10:e02215–22.

42. Olson DK, Yoshizawa S, Boeuf D, Iwasaki W, DeLong EF. 2018. Proteorhodopsin variability and distribution in the North Pacific Subtropical Gyre. The ISME Journal 12:1047–1060.

43. Strauss J, Deng L, Gao S, Toseland A, Bachy C, Zhang C, Kirkham A, Hopes A, Utting R, Joest EF, Tagliabue A, Löw C, Worden AZ, Nagel G, Mock T. 2023. Plastid-localized xanthorhodopsin increases diatom biomass and ecosystem productivity in iron-limited surface oceans. Nat Microbiol 8:2050–2066.

44. Giovannoni SJ. 2017. SAR11 Bacteria: The most abundant plankton in the oceans. Annu Rev Mar Sci 9:231–255.

45. Zhang L, Meng L, Fang Y, Ogata H, Okazaki Y. 2024. Spatiotemporal dynamics of giant viruses within a deep freshwater lake reveal a distinct dark-water community. The ISME Journal 18:wrae182.

46. Pitot TM, Girard C, Rapp JZ, Somerville V, Culley AI, Vincent WF, Moineau S, Roux S. 2025. Viral niche-partitioning: comparative genomics of giant viruses across environmental gradients in a high Arctic freshwater-saltwater lake. ISME Communications 5:ycae155.

47. Fang J, Zhang Y, Zhu T, Li Y. 2023. Scramblase activity of proteorhodopsin confers physiological advantages to Escherichia coli in the absence of light. iScience 26:108551.

48. Ran T, Ozorowski G, Gao Y, Sineshchekov OA, Wang W, Spudich JL, Luecke H. 2013. Cross-protomer interaction with the photoactive site in oligomeric proteorhodopsin complexes. Acta Crystallogr D Biol Crystallogr 69:1965–1980.

49. Needham DM, Poirier C, Hehenberger E, Jiménez V, Swalwell JE, Santoro AE, Worden AZ. 2019. Targeted metagenomic recovery of four divergent viruses reveals shared and distinctive characteristics of giant viruses of marine eukaryotes. Phil Trans R Soc B 374:20190086.

50. Filée J, Siguier P, Chandler M.. 200618. I am what I eat and I eat what I am: acquisition of bacterial genes by giant viruses. Giant viruses and their mobile genetic elements: the molecular symbiosis hypothesis. Trends Genet. Current Opinion in Virology 23(1)1-15.33:81–88.

51. Letelier RM, White AE, Bidigare RR, Barone B, Church MJ, Karl DM. 2017. Light absorption by phytoplankton in the North Pacific Subtropical Gyre. Limnology & Oceanography 62:1526–1540.

52. Keller MD, Selvin RC, Claus W, Guillard RRL. 1987. Media for the culture of oceanic ultraphytoplankton. Journal of Phycology 23:633–638

53. Li Q, Edwards KF, Schvarcz CR, Selph KE, Steward GF. 2021. Plasticity in the grazing ecophysiology of *Florenciella* (Dichtyochophyceae), a mixotrophic nanoflagellate that consumes *Prochlorococcus* and other bacteria. Limnology & Oceanography 66:47–60.

54. Bratbak G, Jacobsen A, Heldal M, Nagasaki K, Thingstad F. 1998. Virus production in Phaeocystis pouchetii and its relation to host cell growth and nutrition. Aquat Microb Ecol 16:1–9.

55. Brussaard CPD. 2004. Optimization of Procedures for Counting Viruses by Flow Cytometry. Appl Environ Microbiol 70:1506–1513.

56. Christaki U, Courties C, Massana R, Catala P, Lebaron P, Gasol JM, Zubkov MV. 2011. Optimized routine flow cytometric enumeration of heterotrophic flagellates using SYBR Green I. Limnology & Ocean Methods 9:329–339.

57. Zhao Y, Zhao Y, Zheng S, Zhao L, Zhang W, Xiao T, Grégori G. 2023. Enhanced resolution of marine viruses with violet side scatter. Cytometry Pt A 103:260–268.

58. Doane FW, Anderson N. 1987. Electron microscopy in diagnostic virology: a practical guide and atlas. Cambridge University Press, New York.

59. Schindelin J, Arganda-Carreras I, Frise E, Kaynig V, Longair M, Pietzsch T, Preibisch S, Rueden C, Saalfeld S, Schmid B, Tinevez J-Y, White DJ, Hartenstein V, Eliceiri K, Tomancak P, Cardona A. 2012. Fiji: an open-source platform for biological-image analysis. Nat Methods 9:676–682.

60. PacificBiosciences. pbdmarkdup. https://github.com/PacificBiosciences/pbmarkdup (25 February 2025, date last accessed)

61. BioInfoTools. BBMap. https://github.com/BioInfoTools/BBMap/blob/master/sh/bbduk.sh (21 February 2025, date last accessed)

62. Kbaseapps. BBTools. https://github.com/kbaseapps/BBTools?tab=readme-ov-file (21 February 2025, date last accessed)

63. Kolmogorov M, Yuan J, Lin Y, Pevzner PA. 2019. Assembly of long, error-prone reads using repeat graphs. Nat Biotechnol 37:540–546.

64. Alneberg J, Bjarnason BS, De Bruijn I, Schirmer M, Quick J, Ijaz UZ, Lahti L, Loman NJ, Andersson AF, Quince C. 2014. Binning metagenomic contigs by coverage and composition. Nat Methods 11:1144–1146.

65. Wu Y-W, Tang Y-H, Tringe SG, Simmons BA, Singer SW. 2014. MaxBin: an automated binning method to recover individual genomes from metagenomes using an expectation-maximization algorithm. Microbiome 2:26.

66. Kang DD, Li F, Kirton E, Thomas A, Egan R, An H, Wang Z. 2019. MetaBAT 2: an adaptive binning algorithm for robust and efficient genome reconstruction from metagenome assemblies. PeerJ 7:e7359.

67. Pan S, Zhao X-M, Coelho LP. 2023. SemiBin2: self-supervised contrastive learning leads to better MAGs for short- and long-read sequencing. Bioinformatics 39:i21–i29.

68. Pitot TM, Brůna T, Schulz F. 2024. Conservative taxonomy and quality assessment of giant virus genomes with GVClass. npj Viruses 2:60.

69. Vaser R, Sović I, Nagarajan N, Šikić M. 2017. Fast and accurate de novo genome assembly from long uncorrected reads. Genome Res 27:737–746.

70. Aramaki T, Blanc-Mathieu R, Endo H, Ohkubo K, Kanehisa M, Goto S, Ogata H. 2020. KofamKOALA: KEGG Ortholog assignment based on profile HMM and adaptive score threshold. Bioinformatics 36:2251–2252.

71. Cantalapiedra CP, Hernández-Plaza A, Letunic I, Bork P, Huerta-Cepas J. 2021. eggNOG-mapper v2: functional annotation, orthology assignments, and domain prediction at the metagenomic scale. Molecular Biology and Evolution 38:5825–5829.

72. Camacho C, Coulouris G, Avagyan V, Ma N, Papadopoulos J, Bealer K, Madden TL. 2009. BLAST+: architecture and applications. BMC Bioinformatics 10:421.

73. Chan PP, Lin BY, Mak AJ, Lowe TM. 2021. tRNAscan-SE 2.0: improved detection and functional classification of transfer RNA genes. Nucleic Acids Research 49:9077–9096.

74. Minh BQ, Schmidt HA, Chernomor O, Schrempf D, Woodhams MD, Von Haeseler A, Lanfear R. 2020. IQ-TREE 2: New models and efficient methods for phylogenetic inference in the genomic era. Molecular Biology and Evolution 37:1530–1534.

75. Benson DA, Cavanaugh M, Clark K, Karsch-Mizrachi I, Lipman DJ, Ostell J, Sayers EW. 2012. GenBank. Nucleic Acids Research 41:D36–D42.

76. Eddy SR. 2011. Accelerated profile HMM searches. PLoS Comput Biol 7:e1002195.

77. Katoh K, Standley DM. 2013. MAFFT multiple sequence alignment software version 7: improvements in performance and usability. Molecular Biology and Evolution 30:772–780.

78. Capella-Gutiérrez S, Silla-Martínez JM, Gabaldón T. 2009. trimAl: a tool for automated alignment trimming in large-scale phylogenetic analyses. Bioinformatics 25:1972–1973.

79. NeLli-team. nsgtree. https://github.com/NeLLi-team/nsgtree (03 March 2024, date last accessed)

80. Letunic I, Bork P. 2024. Interactive Tree of Life (iTOL) v6: recent updates to the phylogenetic tree display and annotation tool. Nucleic Acids Research 52:W78–W82.

81. Yau S, Lauro FM, DeMaere MZ, Brown MV, Thomas T, Raftery MJ, Andrews-Pfannkoch C, Lewis M, Hoffman JM, Gibson JA, Cavicchioli R. 2011. Virophage control of antarctic algal host–virus dynamics. Proc Natl Acad Sci USA 108:6163–6168.

82. Villar E, Vannier T, Vernette C, Lescot M, Cuenca M, Alexandre A, Bachelerie P, Rosnet T, Pelletier E, Sunagawa S, Hingamp P. 2018. The Ocean Gene Atlas: exploring the biogeography of plankton genes online. Nucleic Acids Research 46:W289–W295.

83. Vernette C, Lecubin J, Sánchez P, Tara Oceans Coordinators, Acinas SG, Babin M, Bork P, Boss E, Bowler C, Cochrane G, De Vargas C, Gorsky G, Guidi L, Grimsley N, Hingamp1 P, Iudicone D, Jaillon O, Kandels-Lewis S, Karp-Boss L, Karsenti E, Not F, Ogata H, Poulton N, Pesant S, Sardet C, Speich S, Stemmann L, Sullivan MB, Sunagawa S, Wincker P, Sunagawa S, Delmont TO, Acinas SG, Pelletier E, Hingamp P, Lescot M. 2022. The Ocean Gene Atlas v2.0: online exploration of the biogeography and phylogeny of plankton genes. Nucleic Acids Research 50:W516–W526.

84. Mirdita M, Schütze K, Moriwaki Y, Heo L, Ovchinnikov S, Steinegger M. 2022. ColabFold: making protein folding accessible to all. Nat Methods 19:679–682.

85. The UniProt Consortium. 2017. UniProt: the universal protein knowledgebase. Nucleic Acids Res 45:D158–D169.

86. Waman VP, Bordin N, Lau A, Kandathil S, Wells J, Miller D, Velankar S, Jones DT, Sillitoe I, Orengo C. 2025. CATH v4.4: major expansion of CATH by experimental and predicted structural data. Nucleic Acids Research 53:D348–D355.

87. Van Kempen M, Kim SS, Tumescheit C, Mirdita M, Lee J, Gilchrist CLM, Söding J, Steinegger M. 2024. Fast and accurate protein structure search with Foldseek. Nat Biotechnol 42:243–246.

88. Meng EC, Goddard TD, Pettersen EF, Couch GS, Pearson ZJ, Morris JH, Ferrin TE. 2023. UCSF CHIMERAX: Tools for structure building and analysis. Protein Science 32.

