## Supplementary Information for "Evidence for the acquisition of a proteorhodopsin-like rhodopsin by a chrysophyte-infecting giant virus"

List of contents:

1. Table S1
2. Figure S1
3. Figure S2
4. Figure S3
5. Figure S4
6. Supplemental References

**Table S1.** Noteworthy ChrysoHV genes grouped by function, showing their occurrence (tally), position (locus tag), and NCBI GenBank accession ID. The table includes various auxiliary metabolic genes, methyltransferases, glycotransferases, tRNAs, and ribosomal proteins.

| Potential Function | Tally | Locus Tag | Genbank Accession ID |
|---|---|---|---|
| **Cell Wall Structure** | | | |
| N-acetylmuramoyl-L-alanine amidase | 1 | 1143 | XVM29061.1 |
| **Electron Transport Chain** | | | |
| Cytochrome b5 | 2 | 242, 642 | XVM28163.1, XVM28561.1 |
| Flavinator of succinate dehydrogenase | 1 | 538 | XVM28459.1 |
| Succinate dehydrogenase subunit | 3 | 766, 767, 769 | XVM28684.1, XVM28685.1, XVM28687.1 |
| Prohibitin-like protein | 1 | 927 | XVM28845.1 |
| **Glycosylation** | | | |
| Glycosyltransferase | 15 | 127, 706, 726, 969, 970, 972, 973, 974, 976, 977, 979, 980, 982, 992, 1051 | XVM28048.1, XVM28625.1, XVM28645.1, XVM28887.1, XVM28888.1, XVM28890.1, XVM28891.1, XVM28892.1, XVM28894.1, XVM28895.1, XVM28897.1, XVM28898.1, XVM28900.1, XVM28910.1, XVM28969.1 |
| **Iron/Redox Related** | | | |
| 2OG-Fe(II) oxygenase | 1 | 750 | XVM28668.1 |
| Fe-S biosynthesis | 1 | 775 | XVM28693.1 |
| Fe-containing alcohol dehydrogenase | 1 | 968 | XVM28886.1 |

| | | | |
|---|---|---|---|
| **Light/UV Stress** | | | |
| Photolyase | 3 | 95, 104, 530 | XVM28016.1, XVM28025.1, XVM28451.1 |
| Chlorophyll A-B binding protein | 1 | 850 | XVM28768.1 |
| **Methylation** | | | |
| Methyltransferase | 7 | 44, 113, 160, 684, 741, 988, 1131 | XVM27969.1, XVM28034.1, XVM28081.1, XVM28603.1, XVM28660.1, XVM28906.1, XVM29049.1 |
| **Phosphate Limitation** | | | |
| Phosphate starvation protein (phoH) | 1 | 1007 | XVM28925.1 |
| **Sulfonation** | | | |
| Sulfotransferase | 2 | 1035, 1046 | XVM28953.1, XVM28964.1 |
| **Tail Structure** | | | |
| Tail fiber domain-containing proteins | 2 | 194, 986 | XVM28115.1, XVM28911.1 |
| **Transport** | | | |
| ABC transporter | 7 | 94, 247, 713, 800, 920, 951, 961 | XVM28015.1, XVM28168.1, XVM28632.1, XVM28718.1, XVM28838.1, XVM28869.1, XVM28879.1 |
| Ammonium transporter | 2 | 689, 1074 | XVM28608.1, XVM28992.1 |
| Porins | 1 | 531, 650, 651 | XVM28452.1, XVM28569.1, XVM28570.1 |
| SemiSWEET sugar transporter | 1 | 203 | XVM28124.1 |

| | | | |
|---|---|---|---|
| Urea transporter | 2 | 1087, 1088 | XVM29005.1, XVM29006.1 |
| **Translation** | | | |
| Ribosomal protein S27a | 1 | 1099 | XVM29017.1 |
| tRNAs | 7 | 75, 76, 77, 78, 615, 616, 748 | *NA* |
| Ubiquitin-60S ribosomal protein L40 | 1 | 536 | XVM28457.1 |
| **Rhodopsin** | | | |
| Heliorhodopsin | 2 | 106, 938 | XVM28027.1, XVM28856.1 |
| Proteorhodopsin | 1 | 101 | XVM28022.1 |

31

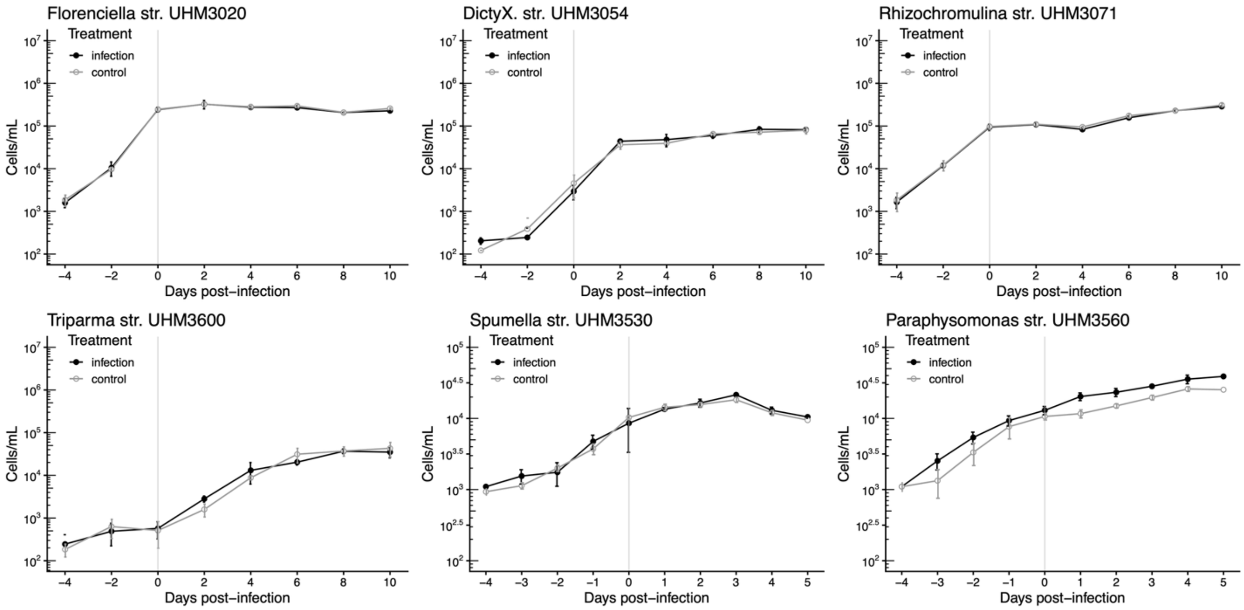


**Fig. S1. Mixotrophic and heterotrophic protists resistant to ChrysoHV.** Time series of protists isolated from Station ALOHA: *Florenciella* str. UHM3020, Dictyochophyceae Clade X str. UHM3054, *Rhizochromulina* str. UHM3071, *Triparma* str. UHM3600, *Spumella* str. UHM3530, and *Paraphysomonas* str. UHM3560, challenged with raw ChrysoHV lysate at time 0. The infection time series is shown in black with filled points, and the control time series is shown in gray with open points.

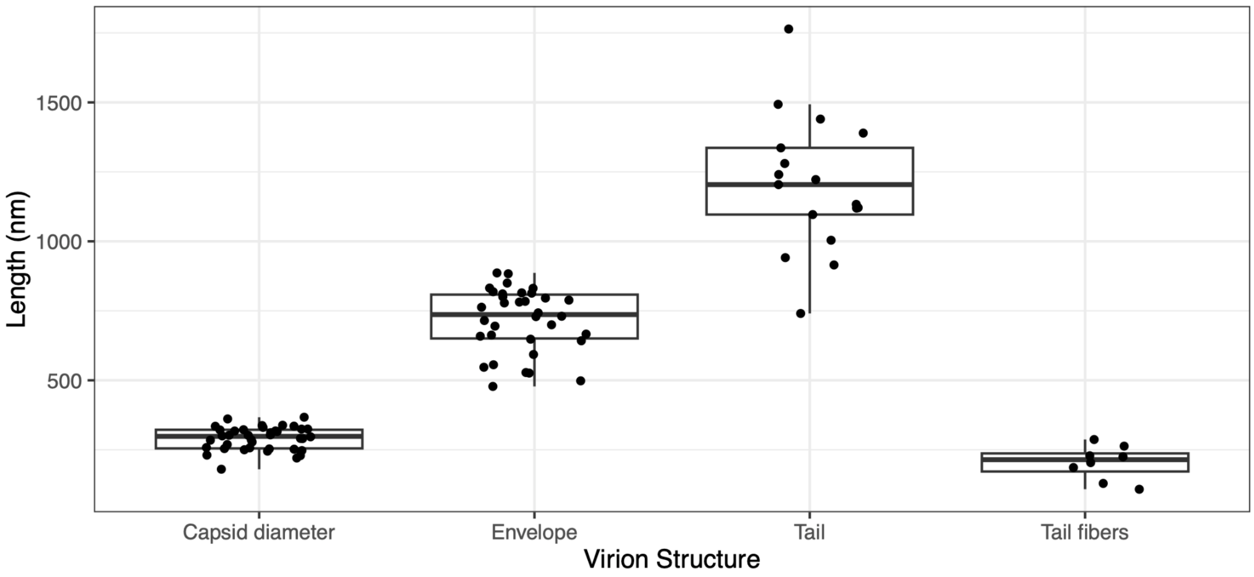


**Fig. S2. Measurements of ChrysoHV virion features.** Each dot represents a data point and each box denotes boundaries of the upper and lower quartiles of each measurement, and the thick, black horizontal line denotes the median value. The whiskers extend out to non-outlier minimum and maximum values.

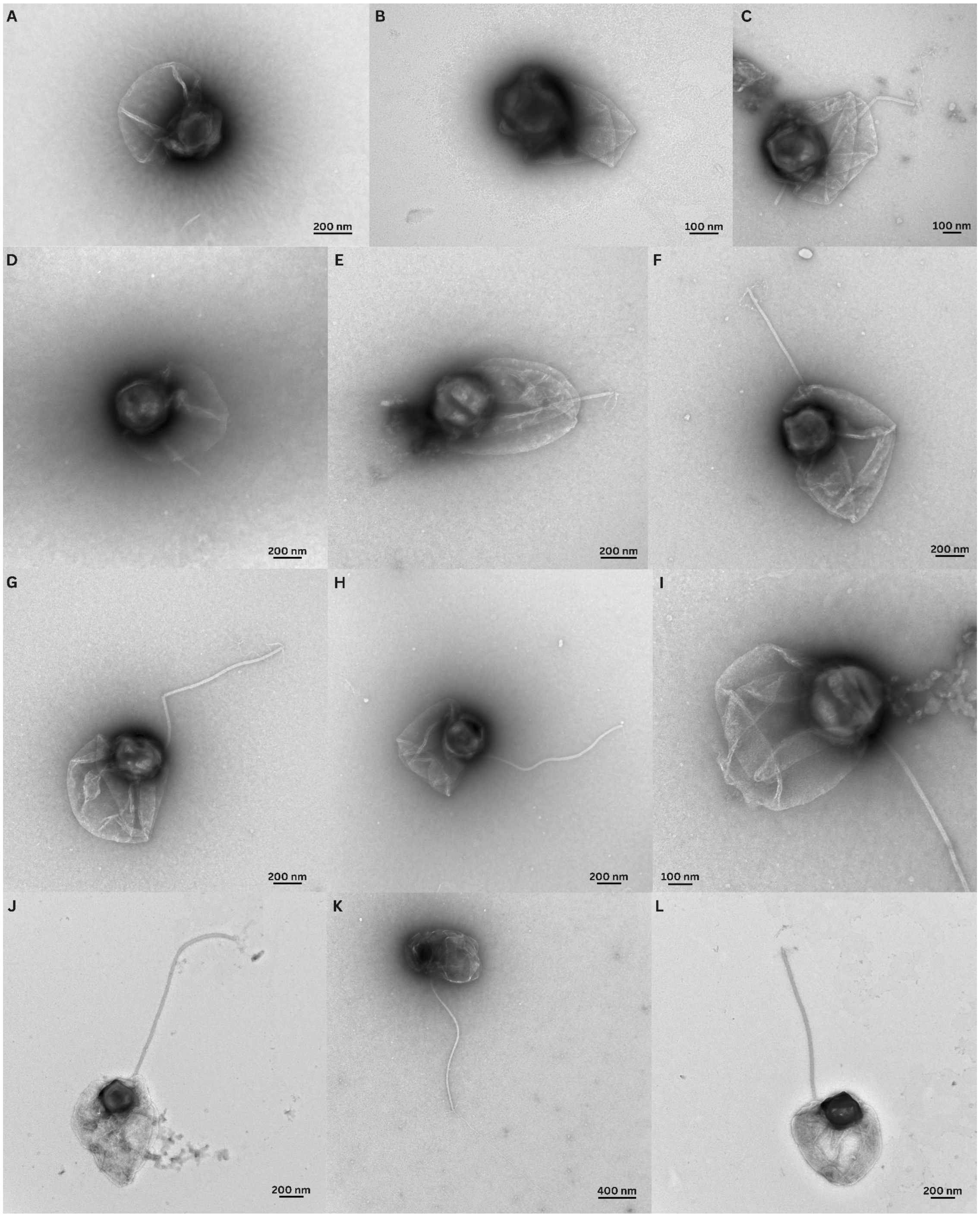


**Fig. S3. TEM micrographs of ChrysoHV.** The images show (A, B) non-tailed morphotypes propagated on chrysophyte Clade H UHM3501; (C, D, E) virions with the tail folded underneath the capsid and envelope (lysate from chrysophyte Clade H UHM3500); and (F-L) tailed morphotypes propagated on UHM3500.

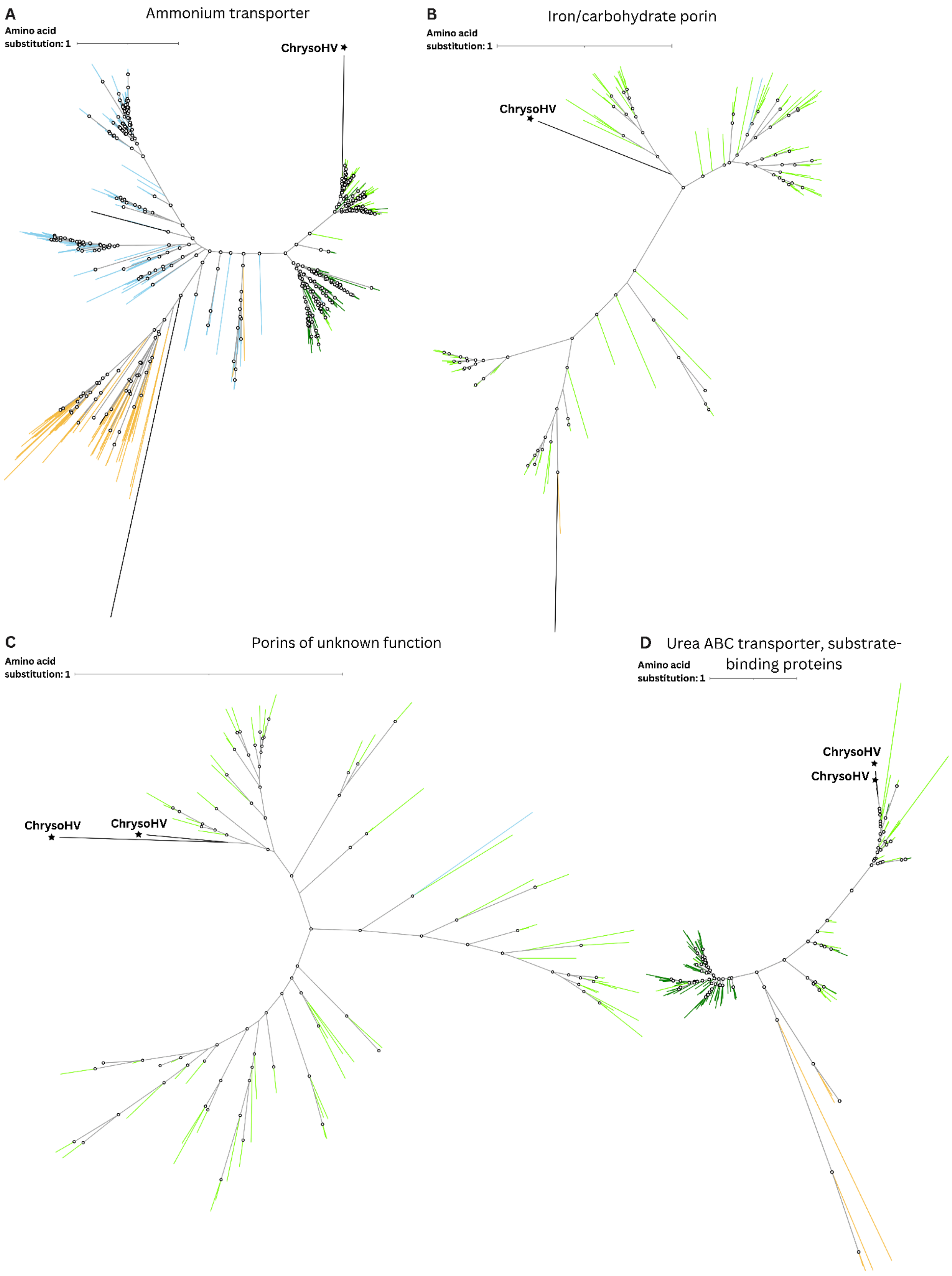


**Fig. S4. (continues on next page)**

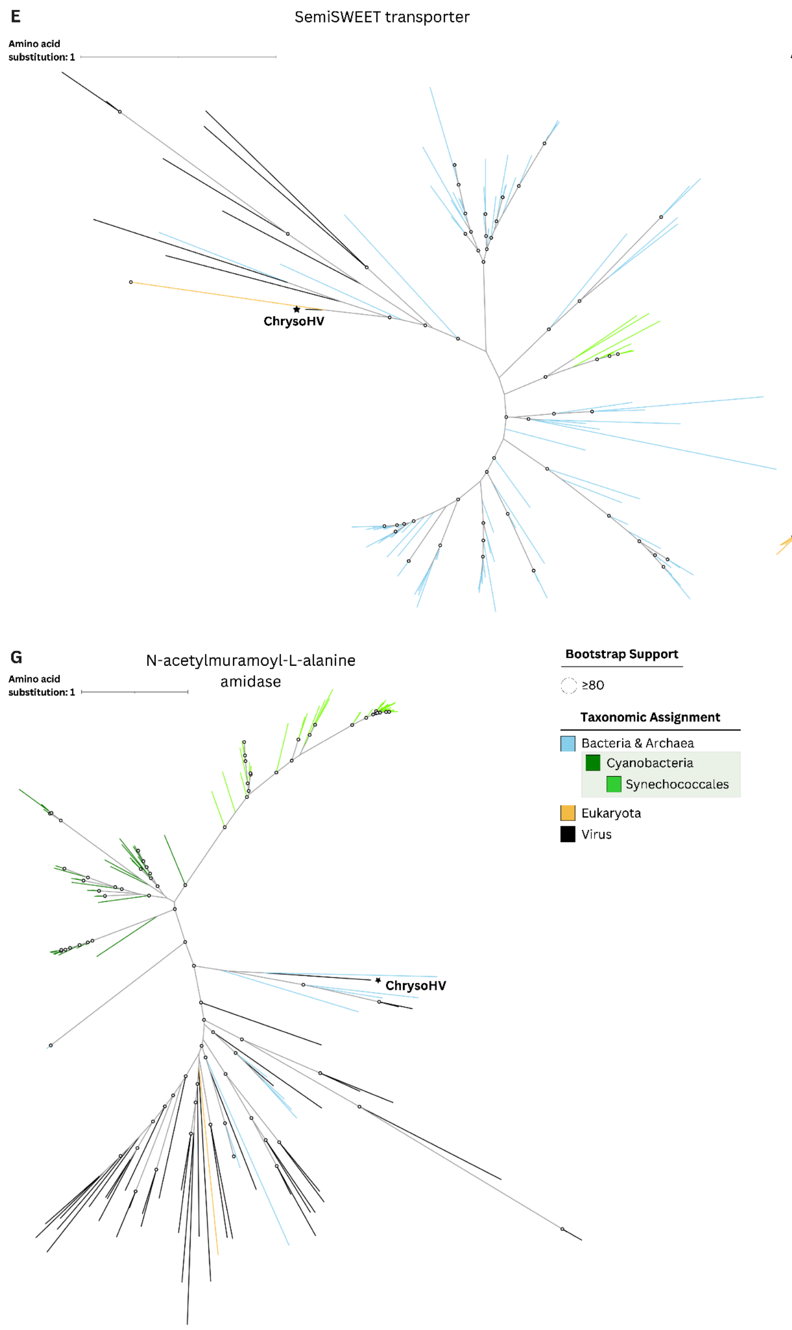


**Fig. S4. Maximum likelihood phylogenies of cyanobacterial homologs found in ChrysoHV.** The gene trees include an (A) ammonium transporter (XVM28992.1), (B) iron/carbohydrate porin (XVM28452.1), (C) porins of unknown function (XVM28569.1, XVM28570.1), (D) urea ABC transporter, substrate-binding proteins (XVM29005.1, XVM29006.1), (E) semiSWEET sugar transporter (XVM28124.1), (F) sulfotransferase (XVM28964.1), and (G) N-acetylmuramoyl-L-alanine amidase (XVM29061.1). The query sequences represent top BLASTp

(1) hits retrieved from the NCBI ClusteredNR database, NCBI Virus database, and Marine Microbial Eukaryote Transcriptome Sequencing Project (MMETSP) (2, 3), then quality filtered for a query coverage ≥ 50% and an E-value ≤ 1e-05. Not all gene trees contained viral or eukaryotic sequences that met these criteria. For closely related genes, such as the porins (C) and urea substrate-binding proteins (D), these top hits were dereplicated and combined into one tree. Hits to the same transcriptome were also dereplicated. The query sequences were aligned with MAFFT (L-INS-i; v7.520) (4), trimmed with trimAl (-gt 0.1; v1.4.1) (5); and the phylogeny was built with IQ-TREE best-fit substitution model and ultrafast bootstrap supports (-m TEST, -bb 1000; v2.2.6) (6), then annotated in iTOL (v6) (7). Branch color indicates sequence taxonomy with the order Synechococcales, which includes *Prochlorococcus* and *Synechococcus*, in bright green; all other cyanobacteria phylum representatives are shown in dark green. Other prokaryotic sequences are shown in light blue, and eukaryote-derived sequences are shown in yellow. Virus sequences are shown in black with ChrysoHV sequences labelled in bold font with a star symbol. Bootstrap support above 80% is indicated by a white circle. Because of their excessive length, branches for sequences YP_010670258.1 and Amphidinium_carterae_Transcript_38023|m.85619 were pruned from the porins of unknown function (C) and urea substrate-binding protein (D) gene trees.

### References


1. Camacho C, Coulouris G, Avagyan V, Ma N, Papadopoulos J, Bealer K, Madden TL. 2009. BLAST+: architecture and applications. *BMC Bioinformatics* 10:421.

2. Keeling PJ, Burki F, Wilcox HM, Allam B, Allen EE, Amaral-Zettler LA, Armbrust EV, Archibald JM, Bharti AK, Bell CJ, Beszteri B, Bidle KD, Cameron CT, Campbell L, Caron DA, Cattolico RA, Collier JL, Coyne K, Davy SK, Deschamps P, Dyhrman ST, Edvardsen B, Gates RD, Gobler CJ, Greenwood SJ, Guida SM, Jacobi JL, Jakobsen KS, James ER, Jenkins B, John U, Johnson MD, Juhl AR, Kamp A, Katz LA, Kiene R, Kudryavtsev A, Leander BS, Lin S, Lovejoy C, Lynn D, Marchetti A, McManus G, Nedelcu AM, Menden-Deuer S, Miceli C, Mock T, Montresor M, Moran MA, Murray S, Nadathur G, Nagai S, Ngam PB, Palenik B, Pawlowski J, Petroni G, Piganeau G, Posewitz MC, Rengefors K, Romano G, Rumpho ME, Rynearson T, Schilling KB, Schroeder DC, Simpson AGB, Slamovits CH, Smith DR, Smith

GJ, Smith SR, Sosik HM, Stief P, Theriot E, Twary SN, Umale PE, Vaulot D, Wawrik B, Wheeler GL, Wilson WH, Xu Y, Zingone A, Worden AZ. 2014. The Marine Microbial Eukaryote Transcriptome Sequencing Project (MMETSP): Illuminating the Functional Diversity of Eukaryotic Life in the Oceans through Transcriptome Sequencing. *PLoS Biol* 12:e1001889.

3. Johnson, Lisa K., Alexander, Harriet, & Brown, C. Titus. (2018). MMETSP re-assemblies. https://doi.org/10.5281/zenodo.740440 (06 July 2026, date last accessed)

4. Katoh K, Standley DM. 2013. MAFFT multiple sequence alignment software version 7: improvements in performance and usability. *Molecular Biology and Evolution* 30:772–780.

5. Capella-Gutiérrez S, Silla-Martínez JM, Gabaldón T. 2009. trimAl: a tool for automated alignment trimming in large-scale phylogenetic analyses. *Bioinformatics* 25:1972–1973.

6. Minh BQ, Schmidt HA, Chernomor O, Schrempf D, Woodhams MD, Von Haeseler A, Lanfear R. 2020. IQ-TREE 2: New models and efficient methods for phylogenetic inference in the genomic era. *Molecular Biology and Evolution* 37:1530–1534.

7. Letunic I, Bork P. 2024. Interactive Tree of Life (iTOL) v6: recent updates to the phylogenetic tree display and annotation tool. *Nucleic Acids Research* 52:W78–W82.
